# Prebiotic xylo-oligosaccharides for alleviation of hepatic steatosis: Results from a four-month dietary intervention and determinants of response

**DOI:** 10.1101/2025.02.19.638992

**Authors:** Jukka E Hintikka, Perttu Permi, Marko Lehtonen, Sanna Lensu, Anastasiia Driuchina, Mika Jormanainen, Jari Laukkanen, Nitin Bayal, Pande P. Erawijantari, Thomas M O’Connell, Leo Lahti, Satu Pekkala

## Abstract

A common complexity from increasing rates of overweight is metabolic dysfunction associated steatotic liver disease (MASLD) which can develop into more severe conditions such as fibrosis or hepatocellular carcinoma. The gut and its microbial flora interact with the liver through the gut-liver axis and supporting healthy composition and metabolism of the microbiota can benefit metabolic health. We have previously shown in animals that a xylo-oligosaccharide (XOS) prebiotic can drive beneficial gut flora and ameliorate hepatic steatosis- our current objective is to validate these results in humans. Forty-two adults (mean age 53.7, mean BMI 33.5) ingested a prebiotic dose of XOS daily for four months. Liver fat was quantified with MRI and body composition was measured. Standard clinical measurements and untargeted metabolomes were analyzed from serum. Microbiota composition, metagenome and metabolites were analyzed from fecal samples. A subgroup analysis was conducted between non-steatotic individuals, non-responders, and responders. The XOS supplementation caused decreases in detrimental amino acid degradation products (AADPs) isovalerate, isobutyrate, and phenylacetate in the gut while serum metabolome remained stable. A decrease in liver fat during the prebiotic was accompanied by a decrease in visceral fat. In a prediction analysis, this response was driven by higher initial ratio of *Bacteroides* to *Faecalibacterium*, higher fecal AADPs, and higher serum amino acids. Non-responders had metabolomic signatures of advanced steatosis, possibly impeding effects from XOS. In overweight individuals with early hepatic steatosis, XOS may correct fermentation imbalances in the gut and decrease production of harmful metabolites, which may benefit hepatic health.

**IMPORTANCE:** Pre- and probiotics are emerging, yet unknown, avenues of treatment for metabolism-associated steatotic liver disease. In overweight humans, a xylo-oligosaccharide (XOS) prebiotic induced shifts in gut bacteria and their metabolites. High proteolytic activity in the gut was predictive of hepatic response to XOS, which may serve the development of personalized treatment of fatty liver.

## INTRODUCTION

Metabolic dysfunction associated steatotic liver disease (MASLD) is a condition characterized by an excessive accumulation of fat in the liver (>5% fat), in absence of excessive alcohol intake or steatogenic drugs. Disproportionately affecting individuals with excess weight, MASLD is present in approximately 75% of people with overweight and in 90% of people with obesity (1). If not properly treated, MASLD can eventually progress to cirrhosis, and even liver cancer (1, 2). Despite the severity, the condition is treatable by adopting healthy lifestyle and addressing adjacent health problems. Fat mass loss of 5-10% can decrease liver steatosis significantly, which is why caloric restriction and prescription of physical activity are the primary non-pharmaceutical avenues of treatment (3). However, since adherence to extensive lifestyle interventions is inconsistent and depends on personal resources (4), auxiliary treatment methods should be explored.

The gut microbiota interacts with the liver through the gut-liver axis, which is a bidirectional communication system that involves metabolic products, immune cells, hormones, and neural pathways (5). Dysbiosis of the gut microbiota has been implicated in the pathogenesis and progression of MASLD. Leakage of bacterial endotoxins and metabolites, such as lipopolysaccharides, into the portal circulation may induce hepatic inflammation and fibrosis, alter bile acid signaling, and modulate host energy balance and lipid homeostasis (6). Perturbations in the composition of the gut microbiota has been observed in patients suffering from MASLD with a further loss of diversity in disease progression (2, 7). The discovery of specific microbial signatures may aid the prediction of MASLD progression and severity (8). We have previously described potential microbiome-related taxonomic and chemical biomarkers as indicators of liver disease (9), however, there is still a lack of comprehensive studies of the gut-liver axis focusing on diet and nutrition.

Advances involving the gut liver-axis can also be viewed from therapeutic perspectives. We have previously shown that administration of *Faecalibacterium prausnitzii*, a butyrate-producing abundant member of the human microbiota, into high-fat fed mice alleviated hepatic steatosis and adipose tissue inflammation (10). Further, we could increase the abundance of *F. prausnitzii* through prebiotic xylo-oligosaccharide (XOS) supplementation, and consequently reduce hepatic steatosis in high-fat fed rats (11). The fecal microbiome-metabolome signatures were also associated with decreased adipose tissue inflammation in these animals (12). Thus, we sought to investigate the effects of prebiotic XOS supplement on liver fat content and metabolic health in human individuals with hepatic steatosis.

In a single group control-period study, adults with overweight or obesity were invited as study participants to consume a XOS supplement daily for four months and their liver fat status, body composition and clinical parameters were measured. This was preceded by a 1-month control period, where participants continued normal daily routine without XOS (**Figure 1**). By studying the gut microbiome composition and metabolism, as well as serum metabolomes, we also aimed to elucidate microbial mechanisms behind the responses to the prebiotic and, conversely, what factors might impede responses. While XOS has been shown to increase insulin sensitivity (13, 14), to our knowledge, this study is the first to study whether XOS intervention can alleviate hepatic steatosis and visceral adiposity in humans.

**Figure 1.**
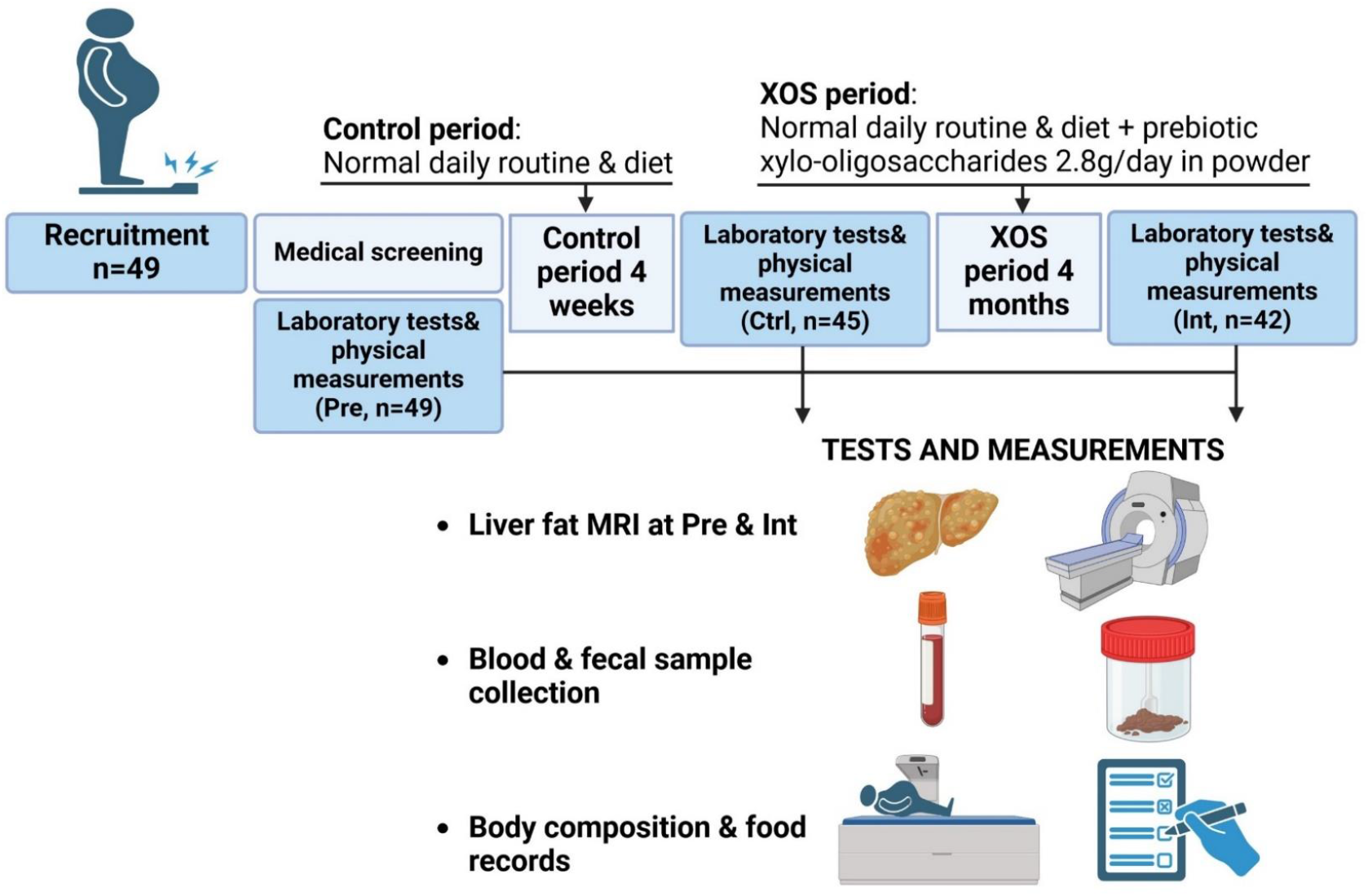
Characteristics of the participants and the study design. The study participants served as their own controls, i.e., we implemented a quasi-experimental design where two time points prior to the intervention were captured. This design has advantages in microbial time-series studies due to the intra-individual variation in the gut microbiome, and also due to the inter-individual variation in the microbiome. Pre = baseline, Ctrl = after 1-month control period, Int = after 4-months of XOS dietary treatment.

## RESULTS

### Biomarkers and body composition showed heterogenous responses during XOS

Forty-nine subjects were enrolled into the study with 42 completing the XOS intervention (**Table 1**). The seven subjects were unable to attend post-intervention measurements due to the onset of COVID-19 and being in the risk groups for infection. Body composition variables showed shifts both during the control period and XOS treatment (**Figure 2**B), suggesting minor lifestyle initiatives by the participants already in the control phase. However, the XOS treatment period was characterized by larger changes in visceral and android fat mass, estimated with bioimpedance and dual-energy X-ray, respectively. Liver enzymes, lipids, and blood glucose levels were analyzed from serum samples using colorimetric methods. Liver fat was quantified using magnetic resonance imaging (MRI) at baseline and after the XOS treatment. Interestingly, very heterogenous responses in liver fat % were observed during the study, with steatosis markedly decreasing in certain individuals and even slightly increasing in others (**Figure 2**A). This prompted us to also conduct subgroup analyses between those who reduced their liver fat by at least 3% (R, responder), those who did not (NR, non-responder), and individuals with low (<5 %) liver fat in baseline (LLF, low liver fat).

**Table 1.**
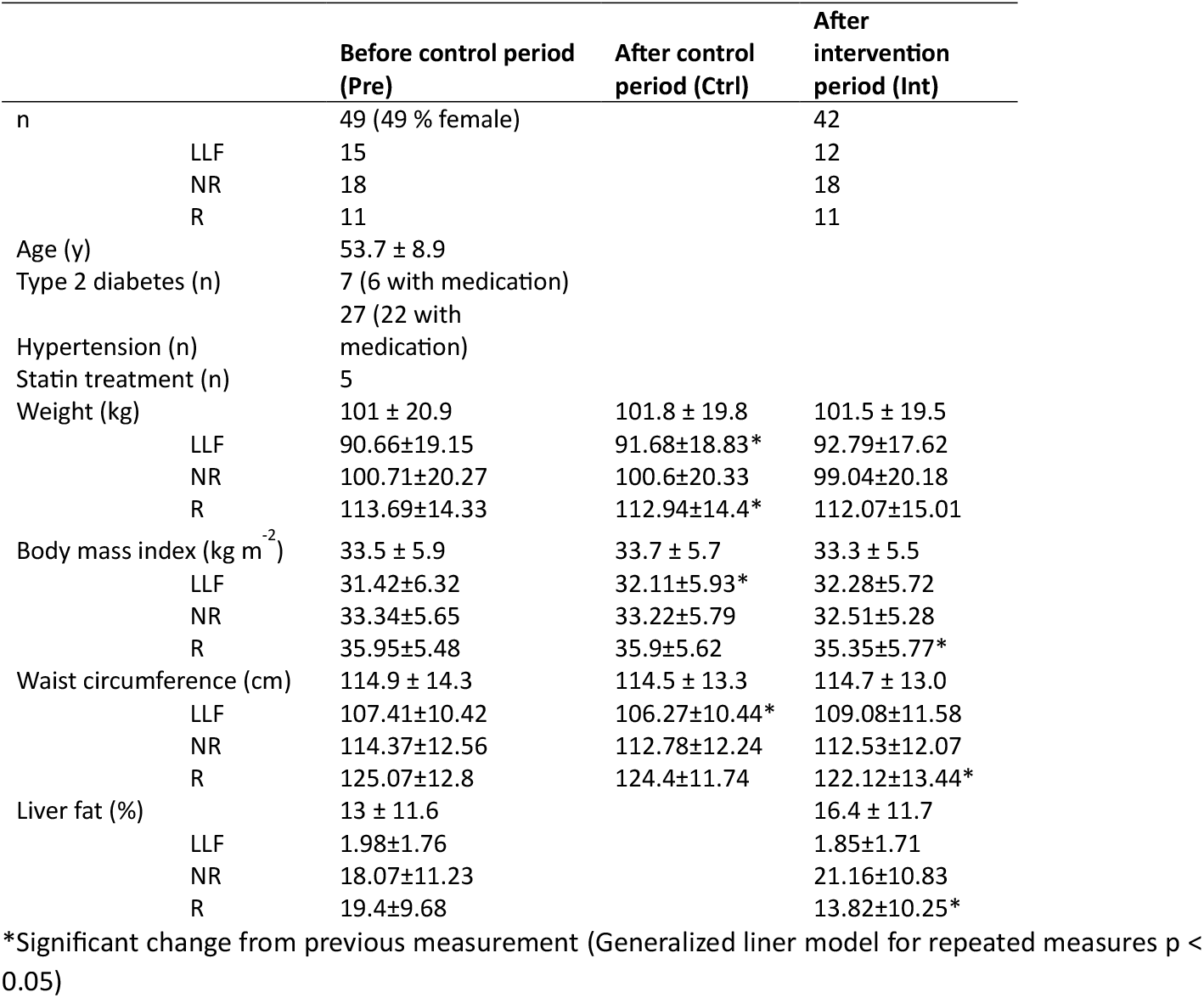
Study sample characteristics for each timepoint and subgroup.

**Figure 2.**
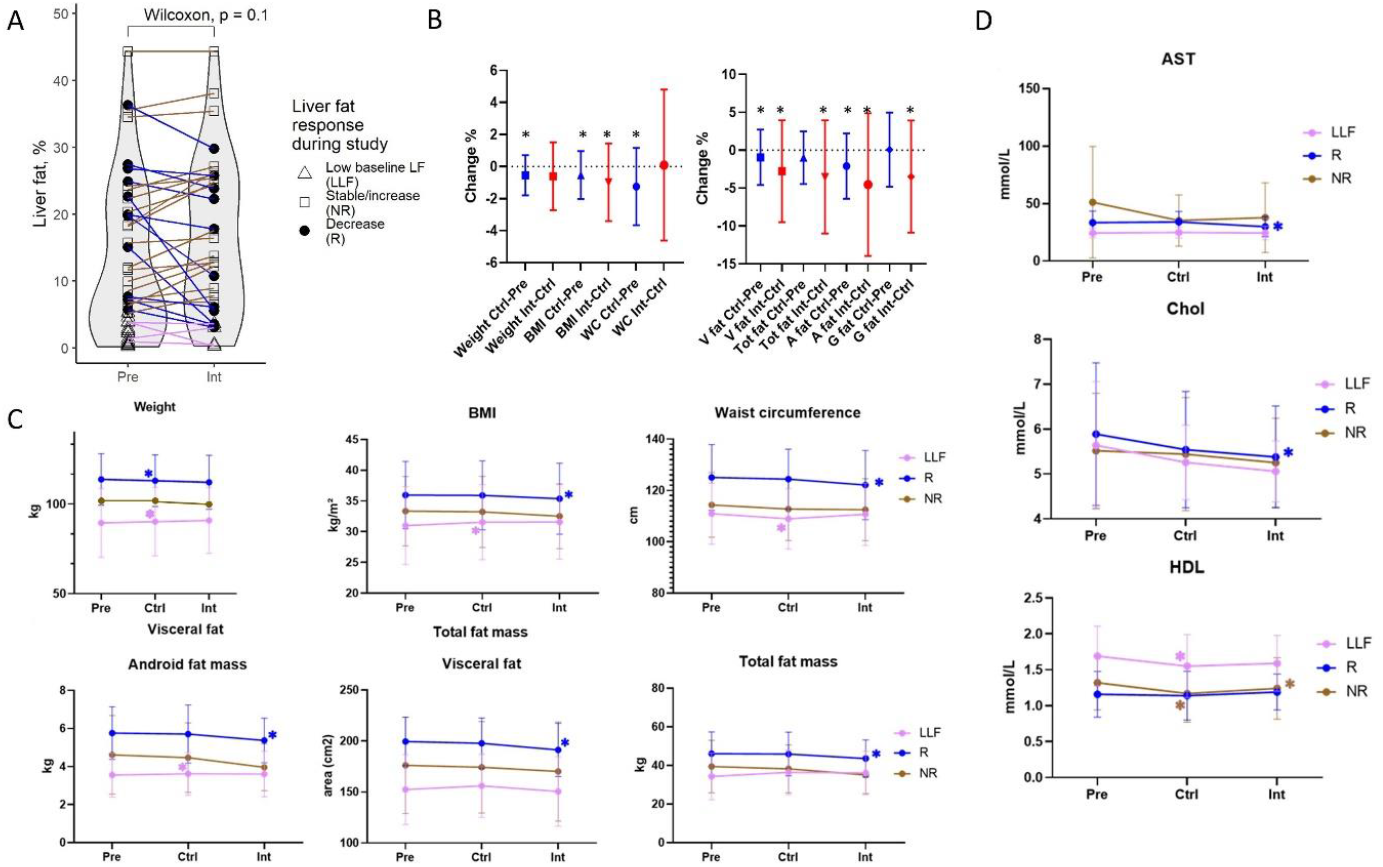
XOS affected liver fat and body composition, but responses were heterogenous. **A)** Liver fat percentage was quantified by MRI. No significant change occurred; thus subgroup analysis was conducted between responders (R, >3% decrease in liver fat), non-responders and low liver fat individuals (<5% at baseline). **B)** Body composition showed slight shifts regardless of study period. Stronger responses in body fat were seen during the XOS period, compared to the control period. **C)** Subgroup analysis of body composition showed that decreases in liver fat were accompanied by decreases in waist circumference, visceral and total fat mass, as apparent by the responders. **D)** Serum cholesterol and HDL slightly improved during the XOS period, with responders also showing improvements in serum AST. Significance tested with Wilcoxon signed rank. * p <0.05. Pre = baseline, Ctrl = after 1-month control period, Int = after 4-months of XOS dietary treatment. LLF = low liver fat, R = responder, NR = non-responder. BMI = body mass index, WC = waist circumference, V fat = visceral fat area, Tot fat = total fat mass, A fat = android fat mass, G fat = gynoid fat mass.

Compared to LLF, the R group was characterized by slightly higher visceral adiposity and fasting glucose both before and after the intervention compared to the other groups (**Supplements, Tables S2 and S3**). Slight changes in body weight were observed during the control period in R and LLF (**Figure 2**C). Within the R group, body mass index (BMI), waist circumference (WC), visceral fat area, total fat mass, android fat mass and gynoid fat mass were found decreased. Of the serum clinical variables, total cholesterol levels decreased during the XOS treatment (**Supplements, Table S1**), while HDL decreased during the control period and increased during the XOS treatment. In addition, XOS treatment slightly improved serum AST and total cholesterol levels in the R group (**Figure 2**D). No changes were detected in insulin, fasting glucose or HOMA-IR during the study.

Diet was recorded in each time point with validated 3-day food diaries (9, 15). Besides a slight decrease in the intake of unsaturated fatty acids in the R group, no changes in diet were observed during the study and the groups were similar in their macronutrient intakes (**Supplements, Table S5**).

### Lactic acid bacteria and proteolytic metabolites decreased during XOS treatment

In 16S rRNA gene sequencing analysis of the microbiota no changes in alpha diversity indices were recorded during the control period or XOS treatment (**Figure 3**B). Based on Bray-Curtis dissimilarity index and PERMANOVA (Permutational Multivariate Analysis of Variance), the gut microbiota beta-diversity did not change between the time points, however the principal coordination plot showed *Faecalibacterium*-dominant communities to diverge from the others (**Figure 3**A). In differential analysis(16), followed by false discovery rate (FDR) correction, the abundances of 13 taxa shifted during the XOS treatment but remained stable during the control period (**Figure 3**C). Among these, we noted increases in Succinivibrionaceae and Gastranaerophilales and decreases in *Ruminococcus gnavus* group and several lactic acid bacteria. Shotgun metagenomic analysis, conducted for R and NR individuals from timepoints Ctrl and Int, did not find changes in microbial taxa, diversity indices, metabolic functions (KEGG orthologies), or Metacyc pathways during the study.

**Figure 3.**
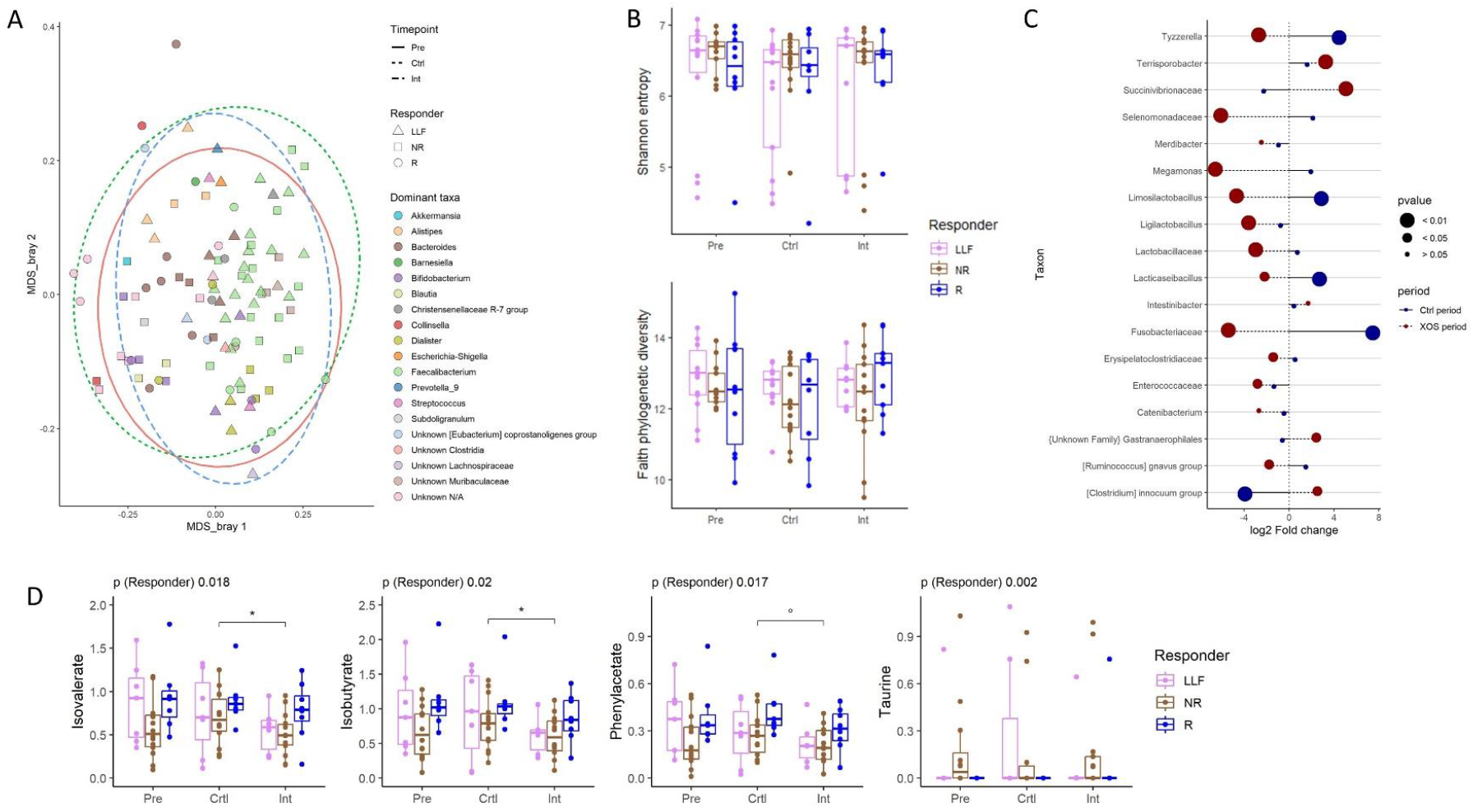
Lactic acid bacteria and proteolytic products decreased during XOS treatment **A)** Bray-Curtis ordination of gut microbial genera showed no longitudinal changes, however the communities showed a separation into Bacteroides-dominant and Faecalibacterium-dominant microbial communities. **B)** OTU-level Shannon diversity and phylogenetic diversity remained stable across the study. **C)** Analysis of the longitudinal changes in gut microbial taxa showed a notable decrease in lactate-producing bacteria during XOS. **D)** Fecal amino acid degradation products (AADPs) isovalerate, isobutyrate, and phenylacetate decreased during XOS treatment and were highest in R. Fecal taurine was lowest in R. Significance tested with mixed linear model. * p <0.05. ° p <0.1 Pre = baseline, Ctrl = after 1-month control period, Int = after 4-months of XOS dietary treatment. LLF = low liver fat, R = responder, NR = non-responder.

Fecal metabolites were analyzed using nuclear magnetic resonance (NMR) spectroscopy. First, NMR spectra were binned into windows of 0.001 ppm and normalized to sum to account for dilution effects. Spectrum-wide changes between timepoints were tested with Spearman correlations, correcting for multiple tests using a method for genome-wide association studies (17). No definite changes were seen during either the control period or the XOS period (**Supplements, Figure S2**). Metabolites were identified using reference spectra, quantified, normalized to probabilistic quotients, and then log-transformed. Statistical analysis was performed on normalized concentrations using linear mixed models with timepoint as a fixed effect and subject as a random effect. Three amino acid degradation products (AADPs) isobutyrate, isovalerate, and phenylacetate were found to decrease in unison during the XOS treatment (**Figure 3**D).

### Responders were characterized by a higher ratio of *Bacteroides* to *Faecalibacterium* and proteolytic metabolites

In group-wise analysis of 16S rRNA data, no longitudinal changes in microbiota diversity indices were detected. Within the time points the LLF, R and NR groups did not differ from each other. At baseline, the phylogenetic diversity of the LLF group tended to be higher than in NR, and the phylogenetic diversity tended to increase in the NR group during the control period **(Supplements, Figure S1)**. Based on Bray Curtis dissimilarity and PERMANOVA, the gut microbiota beta-diversity did not differ between the groups in any time point and no longitudinal changes were detected. The XOS treatment increased the abundance of *Terrisporobacter* and decreased *Lacticaseibacillus* in the R group, and increased *Succinivibrio* in NR (**Supplements, Figure S6**). After the XOS treatment, R also had higher proportions of Desulfobacterota, Anaerovoracaceae, Peptostreptococcaceae and lower proportions of Enterobacteriaceae, Streptococcaceae, Prevotellaceae, Cyanobacteriaceae compared to LLF. (**Figure 4**B). Most notably, the responder group was characterized by an increased ratio of *Bacteroides* to *Faecalibacterium* (BTFR), possibly indicating active proteolytic fermentation in the gut microbiome (18). This difference diminished and lost significance after the XOS treatment (**Figure 4**C). Adjusting for diet did not affect the differences in microbial composition between the responder groups (**Figure 4**A), however the intake of carbohydrates seemed to mediate the group-time interaction in BTFR, as assessed by generalized linear models (**Supplements, Table S6**).

**Figure 4.**
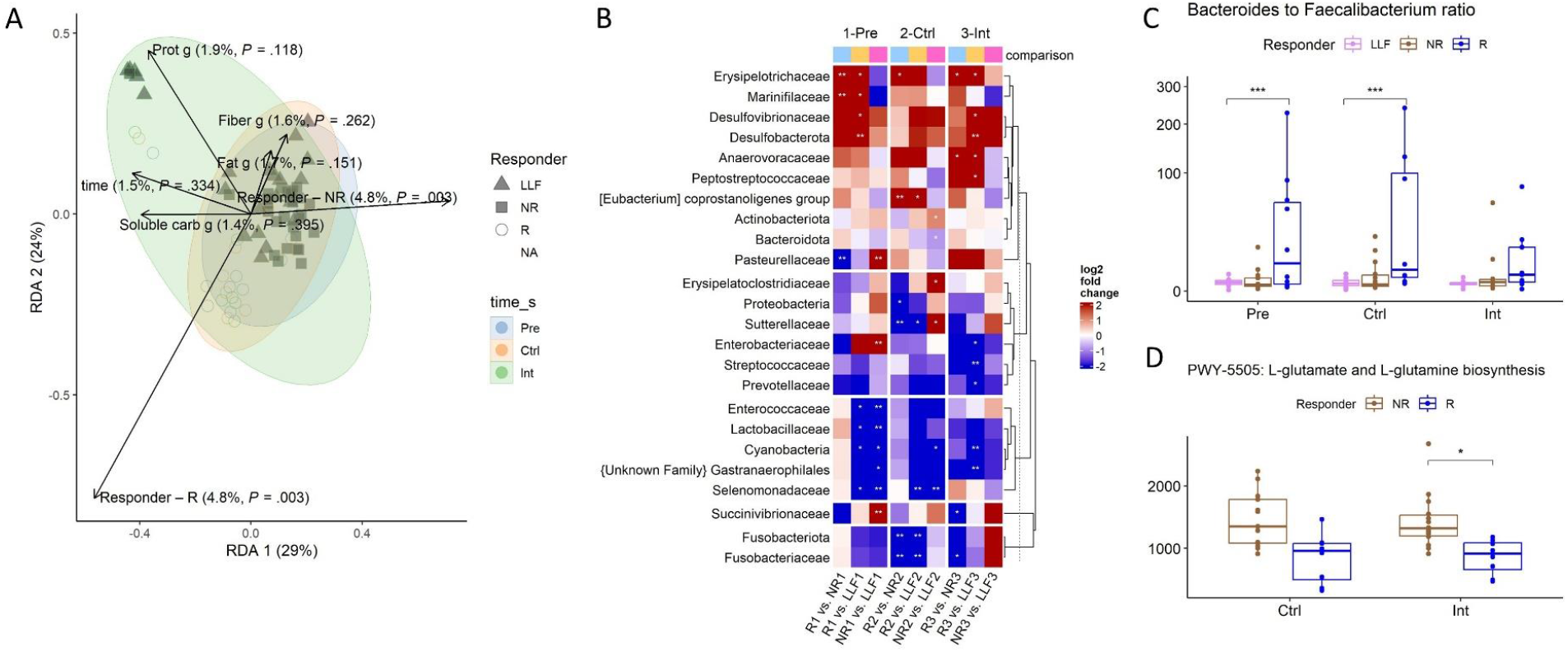
Responders were characterized by a higher ratio of Bacteroides to Faecalibacterium and proteolytic metabolites. **A)** Redundancy analysis (RDA) of the effects of group and diet on gut microbiota. Diet did not affect overall composition or modulate groupwise differences. Significance tested with PERMANOVA. **B)** Differential taxa at family and phylum level in each timepoint. Responders were characterized by elevated abundances of Desulfobacterota and Erysipelotrichaceae. Significance tested with generalized linear model. **C)** Responders had higher ratio of Bacteroides to Faecalibacterium, however, the differences diminished during the XOS treatment. These genera are abundant in the normal human microbiome. Significance tested with Wilcoxon rank-sum. **D)** In the metagenome, the pathway for glutamate and glutamine biosynthesis was more abundant in NR compared to R. Significance tested with Wilcoxon rank-sum. * p <0.05 *** p <0.001 Pre = baseline, Ctrl = after 1-month control period, Int = after 4-months of XOS dietary treatment, LLF = low liver fat, R = responder, NR = non-responder.

Metagenomic analysis did not reveal differences between R and NR groups or longitudinal changes in the microbial community. However, in the analysis of different pathways, the pathway of L-glutamate and L-glutamine biosynthesis (Metacyc PWY-5505) was more abundant in NR compared to R (**Figure 4**D).

Mixed linear models were then used to test effects of time and group on fecal metabolites. Galactose and pyruvate had significant yet minor group-time interactions, but they were attributable to instability and not to either of the treatment periods. However, temporally stable differences between the groups were found in several metabolites: AADPs were higher in R compared to NR and, interestingly, taurine was practically absent in R but detectable in NR (**Figure 3**D).

### Serum metabolomes reflected liver fat and responder status

Untargeted serum metabolomics were conducted using liquid-chromatography/high resolution mass spectrometry. Quality filtered molecular features were log-transformed, pareto-scaled, and then analyzed using principal component analysis (PCA). Timepoint did not cause visible variation in the serum metabolome (**Figure 5**A). The main drivers of variance in the serum metabolome were liver fat %, weight, and HDL cholesterol which showed the strongest associations with the two first components. Loadings for molecular features showed individuals with fatty liver to have higher concentration of glycine-conjugated bile acids and lower concentrations of indolyl compounds. A partial least squares (PLS) model, fitted to classify between timepoints Ctrl and Int, was unable to detect metabolome-wide changes during the treatment. Single features were tested with simple Kruskal-Wallis tests. For features with FDR-corrected p < 0.05, pairwise Wilcoxon tests with Bonferroni correction for multiple comparisons were performed. Only five unannotated molecular features were found to be stable across the control period with significant changes during the XOS period (**Supplements, Table S4**).

**Figure 5.**
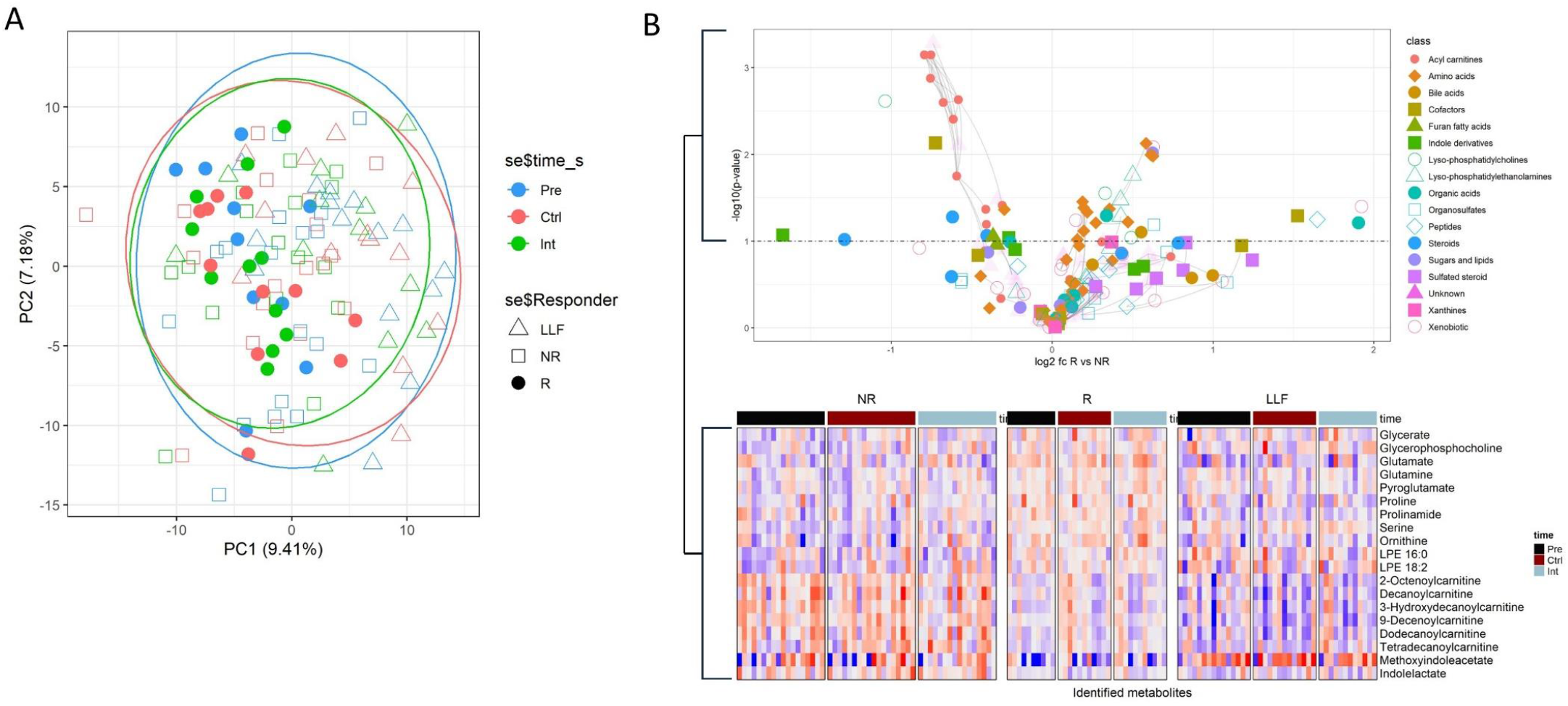
Serum metabolomes reflected liver fat and responder status. **A)** Principal component analysis of the serum metabolome did not show time-wise separation. **B)** Differences in serum metabolome between responder groups. Predictive molecular features of difference between R and NR groups were selected using a sparse partial least squares (PLS) regression and subsequent linear models. Log2 fold change, p-value from linear model, and predicted compound class for each molecular feature shown in the volcano plot. Metabolites identified from the molecular features with p <0.1 visualized with heatmap. Heatmap values are pareto-scaled log2-values. BCAA=branched chain amino acid, LPE=lyso-phosphatidylethanolamine. Pre = baseline, Ctrl = after 1-month control period, Int = after 4-months of XOS dietary treatment, LLF = low liver fat, R = responder, NR = non-responder

Next, the timepoints were combined and used for a pooled analysis to find temporally stable differences in the metabolome between responder groups. The top features from a Sparse PLSDA (VIP >= 1) model were tested with linear models and two-way analyses (responder*time + sex). Of the three groups, responders had the highest levels of glycerate and several amino acids corresponding to enriched glycine-serine metabolism, urea cycle and ammonium recycling (**Figure 5**B). Responders also had higher levels of compounds containing sulfate groups (HO4S^-^), particularly bile acid and other steroids. Medium chain acyl carnitines (C8-C14) decreased groupwise in the order of NR > R > LLF. The lyso-phosphatidylethanolamine species LPE 16:0 and LPE 18:2 behaved in an inverse fashion. The tryptophan metabolite methoxy-indoleacetate showed the largest variation between the groups in the order of LLF > NR > R. Overall, the differences were stable across the study, as assessed by the PLSDA components and the interaction coefficients in the linear models.

### XOS response was predicted by markers of high proteolytic fermentation

Co-varying clusters of metabolites and microbes were analyzed using Pearson correlation, network exploration in Viime(19), and biclustering as done before (12). A group of genera, including *Bacteroides, UCG-002* and *Alistipes* were identified as a potential phenylacetate producers, associating directly with AADPs and inversely with aromatic and branched chain amino acids and Taurine (**Figure 6**A). Short-chain fatty acids and xylose, the monosaccharide unit of XOS, on the other hand associated with a *Faecalibacterium*-driven group of genera. This group of bacteria inversely associated with the AADP cluster, possibly indicating competing fermentation processes of peptides and carbohydrates.

**Figure 6.**
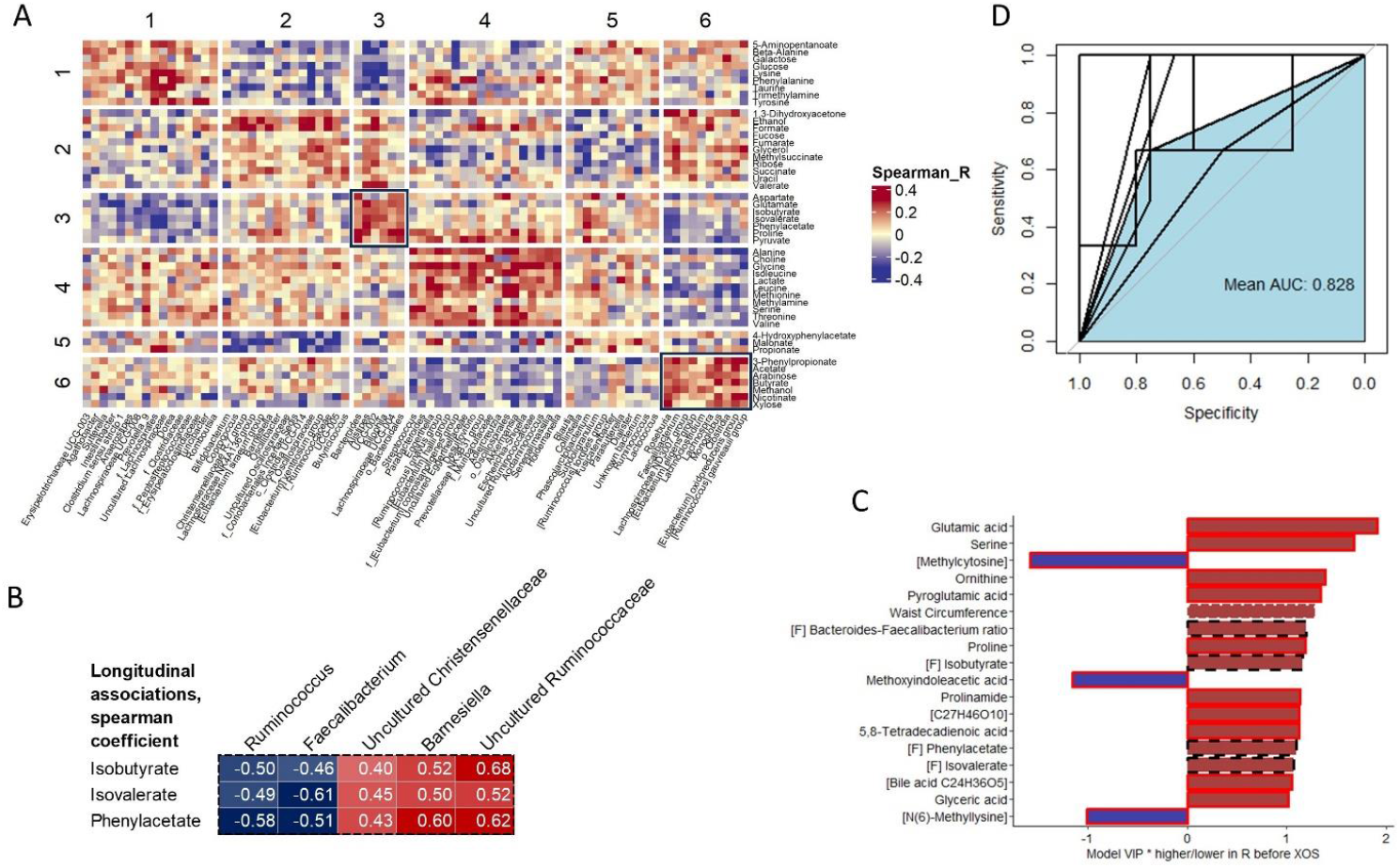
XOS response was predicted by markers of high proteolytic fermentation. **A)** Biclustering of the associations between the gut microbiota genera and fecal metabolites uncovered inversely associated groups of amino acid and carbohydrate degraders **B)** Longitudinal associations were run for changes during the XOS period and showed the decrease in AADPs to associate with Ruminococcus and Faecalibacterium. **C)** Top variables from an OPLS-DA model trained to discern between non-responders and responders before the XOS intervention. The best predictors, besides waist circumference, were signatures of increased protein degradation in the gut and altered circulating amino acids. Black dashed outline = markers in feces. Red solid outline = markers in serum. **D)** ROC plot of the predictive power of the top variables, assessed with 10 fold cross validation. The model reached acceptable performance with an average AUC of 0.828. AUC=area under curve, R= responder

We also analyzed the longitudinal associations between metabolites and microbes by running Spearman correlations on the changes during the XOS period (**Figure 6**B). Interestingly, the decrease in the AADPs associated inversely with changes in the butyrate-producing genera *Faecalibacterium* and *Ruminococcus* and directly with changes in *Barnesiella*, uncultured *Christensenellaceae*, and uncultured *Ruminococcaceae*.

Responsiveness to XOS was investigated by a single nucleotide polymorphism (SNP) analysis Cross reference of SNPs present in the SNP array used and a list of SNPs significantly associated with MASLD in FinnGen’s Freeze(20) 10 identified 13 usable SNPs. Genetic risk for MASLD as calculated four different ways was not significantly correlated with the observed change in liver fat. Association test of selected SNPs with responsiveness did not identify significant associations after multiple test correction, likely due to limited sample size. The most significant SNP, rs3808607 (p-value of 0.0046 before multiple test correction and 0.07 after) was found around 2 kbp upstream of the CYP7A1 gene, suggesting a regulatory function in its expression. While this SNP is marked as significantly associated with MASLD in FinnGen study data Freeze 10 (20), it is listed as benign in the dbSNP database, thus, its relevance to either MASLD or the responsiveness to the prebiotic is unknown.

Finally, we combined the Pre and int timepoints and ran predictive analyses across the omic datasets for response in liver fat % (R vs NR). Features with VIP >1 were selected using O-PLS-DA and tested with logistic regression. With ten cross-validations the model reached acceptable predictive performance with an area under curve (AUC) of 0.828. Serum amino acids and amino acid derivatives were the best predictive variables for the XOS responsiveness, as were waist circumference, fecal AADPs and BTFR (**Figure 6**C-D).

## DISCUSSION

The human responses to probiotics and prebiotics can be highly personal (21, 22) and elucidation of these mechanisms can benefit personalized nutrition for metabolic conditions. As the prevalence of obesity-related comorbidities increases, accessible and cost-effective interventions, such as therapies targeting the gut microbiota, become essential. The results from our previous studies (11, 12) as well as the works of other groups (14, 23, 24) demonstrate a potential in prebiotic XOS supplementation to improve obesity-related conditions of hepatic steatosis, adipose tissue inflammation and insulin resistance. Based on our present and previous studies (10, 11, 25), any potential effects of XOS might be driven by the selective growth of saccharolytic bacteria over certain proteolytic bacteria and the reduced production of detrimental amino acid derivatives in the gut. Responses to prebiotics may be heterogenous, however our study underlines a microbial component in the decrease of hepatic and visceral fat. Individuals with visceral adiposity and high initial proteolytic activity in the gut showed the largest decreases in hepatic fat during the XOS period. Conversely, the serum metabolomes of non-responsive individuals suggested an impaired fatty acid oxidative capacity and advanced inflammation in the liver, a finding that warrants further studies. In the future, our results may serve the development of personalized treatment of MASLD that could be combined with other lifestyle based approaches to improve prognosis and treatment response. However, more research, particularly using individualized interventions, is needed to verify the efficacy of XOS in treatment of metabolic disorders. More research is also needed on how prebiotics work along different dietary patterns.

In this study, co-occurring decreases were seen in fecal phenylacetate, isobutyrate, and isovalerate, which are known markers of hepatic steatosis (26, 27). The XOS-induced decrease in isovalerate in particular corroborates our previous findings in rats (11). The gut microbiome plays a key role in the breakdown of complex dietary components, including proteins that give rise to amino acids. Phenylacetate is produced by the gut microbiota in reactions that degrade aromatic compounds, mainly phenylalanine (28). Similarly, isobutyrate and isovalerate have been linked to degradation of valine and leucine by the gut microbiota(29). Dietary fiber is metabolized by the gut microbiota to beneficial products, mainly short-chain fatty acids that can influence satiety and various physiological processes, directly and through the secretion of certain gut peptides (30–32). Fibers can also reduce the production of detrimental proteolytic metabolites secreted by the gut microbiota which leads to a shift in the overall fermentation balance towards a more beneficial saccharolytic fermentation (30).

Notably, the hepatic response to XOS was accompanied by a high initial BTFR. The inverse of this marker has been linked with hepatic steatosis in a comparable population (25). In the R group, this ratio decreased after the XOS supplementation. *Bacteroides* are not inherently harmful but are known to effectively degrade a wide variety of substrates, including peptides, and this may contribute to the production of detrimental metabolites (33, 34). XOS, on the other hand, is a preferred substrate to *F. prausnitzii*, increasing its *in-vitro growth*(11). Thus, we suspected this shift from high initial peptide fermentation in the gut towards polysaccharide fermentation to be a major mechanism through which XOS might benefit liver health. Interestingly, our results showed that this ratio is an important determinant of the responsiveness to XOS treatment. Hopefully, our findings can have diagnostic utility in the future.

Predictive characteristics for XOS responsiveness before the beginning of the intervention were explored using feature selection and subsequent logistic regression. Higher AADPs and BTFR were among the best predictors, indicating a more active proteolytic fermentation in the gut before the prebiotic supplementation. In addition, the responders were mostly male, had higher waist circumference compared to the other participants and had higher levels of serum amino acids glutamate, serine, ornithine, and proline as well as glycerate. This phenotype and metabotype could be used for selection of responsive participants for personalized interventions, however, larger scale studies with randomized design accompanied with targeted metabolomics should be used to confirm these results. Serum glutamate also exhibited a mild negative correlation (Spearman r -0.37) with the glutamate synthesis pathway of the gut metagenome. Considering this pathway was less abundant in R, a downregulating mechanism would be another future topic of study.

Conversely, the non-responsive individuals had elevated serum concentrations of medium-chain acyl carnitines and decreased concentrations of phosphatidylethanolamine lysoPE16 and lysoPE18. As these are both markers of fatty liver (35, 36), one should consider whether a response to dietary intervention such as XOS could be stunted by advanced fibrosis in the liver. The predictive characteristics described in this study should be validated by replicated studies.

The primary limitation of our study is the limited sample size, and the limited applicability of the results. The direction of causality in the effects seen during XOS supplementation leaves room for interpretation. There is a chance that the reduction in liver fat in R was a result of the slightly reduced visceral adiposity, although our previous studies largely corroborate our conclusions. The main strengths in the study were the use of state-of-the-art methods for omic analyses, the use of a robust, reproducible measurement for liver fat (MRI), and the analysis of the dietary components in each timepoint. Instead of a randomized controlled design, we implemented a quasi-experimental design where two time points prior to the intervention were captured. Despite the limitations, this design has advantages in microbial time-series studies due to the intra-individual variation in the gut microbiome (37) and it also allows to observe regression to the mean in the metabolomes (38)

## MATERIALS AND METHODS

### Study participants

Detailed descriptions of the methods are provided in the **Supplemental data**. The current study was conducted in accordance with the Helsinki Declaration and approved by the Ethics Committee of the Hospital District of Southwest Finland (currently Wellbeing Services County of Southwest Finland) (ETMK 72/2019). A written informed consent was collected from all participants prior to the study.

A total of 49 participants were recruited by the University of Jyväskylä, by advertisements in local bulletins, university’s web pages and social media. The inclusion criteria were an age <75 years, being overweight (body mass index > 25), and high waist circumference (>102 cm for males, >88 cm for females). The exclusion criteria were antibiotic treatment 1 month prior to the study, excessive alcohol consumption (>20 g/day for females, 30 g/day for males), inflammatory bowel disease, celiac disease, major eating disorders, and hypothyroidism. Due to heterogeneity in microbiome-related outcomes, the optimal sample size was determined as the highest possible within the constraints of study resources. The study is registered at ISRCTN under the number ISRCTN86495943.

### Intervention

Each subject underwent a one-month control period before the commencement of XOS prebiotic. During the intervention, the participants ingested 2.8 grams of XOS powder daily for 4 months, dissolving the powder in liquid or yogurt in their normal diet. The dose was determined by previous literature as sufficient to induce change in human gut microbiota (39). During both the control and the XOS period, participants were instructed to resume their normal diet and physical activity. XOS (purity 95%) was purchased in neutral sachets from Van Wankum Ingredients, Maarssen, Netherlands.

### Liver fat quantification

Liver fat was initially screened with ultrasonography. The accurate amount of liver fat was measured with magnetic resonance imaging using the most common technique, out-of-phase/in-phase chemical shift imaging, using two replicate images per measurement (9). The high liver fat (>5% fat) participants were further divided to treatment responders and non-responders based on the decrease/no decrease in liver fat %. Responsiveness was determined as a decrease of at least 3% of liver fat between the two independent image analyses.

### Body composition, clinical variables, and diet

Body composition was assessed using dual-energy X-ray and bioimpedance analyses. The analysis of serum clinical variables was performed using the standard chemistry for liver enzymes, cholesterol, free fatty acids, triglycerides, and glucose. Immunochemistry was used for insulin analysis. Homeostasis model assessment of insulin resistance (HOMA-IR) was calculated as {fasting glucose} * {fasting insulin}/22.5. The diet was analyzed from 3-day self-reported food diaries using Finelli database and food diary analysis web tool (https://fineli.fi). Fecal sampling and food diary filling were performed within the same week.

### Sample collection and handling

Blood samples were collected after a 10 h fasting period by drawing into 10 ml coagulation activator tubes (Vacutest 11030). Serum was separated by centrifugation at 2000 x g for 10 minutes and stored in microcentrifuge tubes at -80° C until analysis. The participants self-collected the fecal samples, which were frozen at domestic freezers immediately after collection, brought to laboratory frozen and stored immediately at −80°C. Total DNA extraction from the fecal samples was performed as described previously (9). Briefly, the DNA was extracted from ∼80 mg of feces with Stool Extraction Kit and semi-automated GenoXtract instrument (Hain Lifescience GmbH, Nehren, Germany), combined with preceding homogenization with bead-beating in 0.5/1.0 mm Ceramic Bead Tubes (Zymo Research, Irvine, CA, USA).

### Gut microbiome

The composition of the gut microbiome was analyzed using 16S rRNA sequencing (IonTorrent, Thermo Fisher Scientific) as previously (9). 16S rRNA gene sequences were clustered into operational taxonomic units (OTUs) at 97% similarity using CLC Microbial Genomics Package (Qiagen, Hilden, Germany). The rRNA gene sequences were classified using SILVA SSU Reference database (v138, 99%).

The samples from individuals with liver fat > 5% in timepoints Ctrl and Int were further subjected to metagenome analysis. Shotgun metagenome sequences were obtained at Novogene (Cambridge, UK). The nextflow pipeline taxprofiler(40) was utilized to preprocess raw reads and taxonomic annotation with MetaPhlan4. The HUMAnN pipeline(41) was utilized to annotate functional data and pathway mapping.

### Metabolomics analyses

For the extraction of fecal metabolites, thawed samples were suspended in phosphate-buffered saline. Fecal metabolites were measured using NMR and quantified with the Chenomx software. For the extraction of serum metabolites, thawed samples were mixed with cold CAN and the compounds were extracted on filter plates. Untargeted metabolomics analysis was conducted on a liquid chromatography-tandem mass spectrometry platform using two different columns and both ionization modes.

### Statistical analyses

Statistical analyses of body composition, clinical variables and diet were performed using IBM SPSS Statistics version 28 (Chicago, IL, USA). Normal distribution of the variables was analyzed using Shapiro Wilk’s test, skewness and curtosis. Statistical analysis of the gut microbial taxa was conducted in CLC microbial genomics package. Metagenomes and metabolomes were analyzed in R (V4.2) Statistical significance was set at P <0.05 unless stated otherwise.

### SNP and predictive analysis

Genotyping was performed by the Institute for Molecular Medicine Finland FIMM Technology Centre, University of Helsinki. A genome wide genotype analysis was made with the customized Illumina Infinium Global Screening Array FIN beadChip (GSAFIN) Sample statistics were calculated using plink (v 1.9). Association of SNPs and MASLD status as well as treatment responder status was calculated via fisher exact test with Lancaster’s mid-p adjustment.

Biclustering of associations between microbial genera and fecal metabolites was performed using an R implementation of spectral coclustering, as proposed by Dhillon (42). For prediction of XOS response, groups R and NR and timepoints Pre and Int were pooled, and best features were selected from an orthogonal partial least squares discriminant analysis (R, package ropls). Logistic regression with 10 cross validations was then run to assess the predictive power of the variables.

## ACKNOWLEDGEMENTS

We would like to thank the laboratory staff of the Faculty of Sport and Health Sciences in University of Jyväskylä for their work in the collection and handling of samples: Mervi Matero, Hanne Tähti, Bettina Hutz and Susanna Luoma. We thank Eeva-Maija Palonen for administrative roles in the project. The authors thank Biocenter Finland and Biocenter Kuopio for supporting the mass spectrometry facilities.

The study was funded by the Research Council of Finland (grant numbers 308042 and 349264), and by the ERVA funding of the Southwest Finland Health Care District. The study was also supported by the Research Council of Finland funded Profiling of University of Jyväskylä and the Faculty of Sport and Health Sciences: Physical activity through life span (PACTS2).

## AUTHOR CONTRIBUTIONS (CREDIT AUTHOR STATEMENT)

JEH: Writing - Original Draft, Visualization, Software PP: Resources, Methodology, Investigation, Supervision ML: Resources, Methodology, Investigation SL: Investigation, Project administration AD: Investigation, Project administration MJ: Formal Analysis JL: Supervision, Project administration NB: Formal analysis PE: Formal analysis TMOC: Methodology, Conceptualization LL: Resources, Methodology, Conceptualization SP: Writing - Original Draft, Conceptualization, Formal Analysis, Funding acquisition, Supervision.

SP and AD had unrestricted access to the research data. All authors contributed to the review and editing of the manuscript and have accepted the final draft.

## DATA AVAILABILITY STATEMENT

The access to the data is restricted due to personal information protection (General Data Protection Regulation (GDPR) 2016/679 and Directive 95/46/EC). However, it is possible to contact author to ask for a copy of the material. The metadata of the study can be found in https://doi.org/10.17011/jyx/dataset/85068.

